# Convolving Pre-Trained Convolutional Neural Networks at Various Magnifications to Extract Diagnostic Features for Digital Pathology

**DOI:** 10.1101/333773

**Authors:** John-William Sidhom, Alexander S. Baras

## Abstract

Deep learning is an area of artificial intelligence that has received much attention in the past few years due to both an increase in computational power with the increased use of graphics processing units (GPU’s) for computational analyses and the performance of these class of algorithms on visual recognition tasks. They have found utility in applications ranging from image search to facial recognition for security and social media purposes. Their continued success has propelled their use across many new domains including the medical field, in areas of radiology and pathology in particular, as these fields are thought to be driven by visual recognition tasks. In this paper, we present an application of deep learning, termed ‘transfer learning’, using ResNet50, a pre-trained convolutional neural network (CNN) to act as a ‘feature-detector’ at various magnifications to identify low and high level features in digital pathology images of various breast lesions for the purpose of classifying them correctly into the labels of normal, benign, in-situ, or invasive carcinoma as provided in the ICIAR 2018 Breast Cancer Histology Challenge (BACH).

## Introduction

While artificial intelligence and machine learning have revolutionized many scientific fields, perhaps no other method has had the widespread adoption and practical use as much as deep artificial neural networks, or otherwise known as ‘deep learning.’ Of note, deep learning has had a profound impact on tasks associated with visual recognition, bringing about technologies capable of object classification and image recognition.^1^ With the reduction to practice in many fields such as social media and communication technologies, there has been a recent advent to bring deep learning into the medical field. One of the initial applications of deep learning in the medical field was by a group at Stanford led by Sebastian Thrun who used transfer learning to classify skin lesions into benign, non-neoplastic, and malignant subtypes.^2^ This proof of concept has led an initiative to bring deep learning into other visual recognition tasks in the medical field including radiology and digital pathology.^3,4^ Not only have these approaches been shown effective in providing diagnostic power, comparable to medical professionals, but they also have shown promise to help learn features possibly missed by humans that can help differentiate various pathologies.^5^

Despite the promise of deep learning for applications in the medical field, due to the novelty of these applications, there exists little labeled data for supervised machine learning. However, prior work in the fields of deep learning has shown the power of a technique called ‘transfer learning,’ the idea being that pre-existing designed architectures (i.e. ResNet50, AlexNet, VGG16) that have been trained on possibly millions of images for over 1000 image classes can serve as ‘professional feature detectors’ for new visual recognition tasks where data may not be as abundant. While it may seem far fetched that a convolutional neural network (CNN) trained to recognize dogs and cats could recognize relevant features in medical imaging, Sebastian Thrun and his group demonstrated this exact concept could be utilized, generating results even better than when training a CNN de-novo to diagnose skin lesions.2 In this manuscript, we propose a method by which one can use ResNet50, a pre-trained CNN, as a feature detector for classification of normal, benign, in-situ, and invasive breast carcinoma. We propose a method by which convolving this pre-trained net at various magnifications of labeled pathology image tiles can serve to detect low and high-level features in digital pathology that can be ultimately used for the task of lesion classification.

## Methodology

### Data-Set

For the task of classifying various types of breast pathology images, we were provided a set of 400 images (100 per class) of normal, benign, in-situ, and invasive breast carcinoma. Additionally, with a set of 20 whole slide images (WSI), we were able to extract an additional 648 image tiles of invasive, in-situ, and normal breast tissue with the assistance of a pathologist. For the remaining of the manuscript, we will refer to the first set as Data Set A and the latter set as Data Set B.

### Color Normalization & Image Augmentation

In order to account for the variation in color we conducted an image color augmentation routine in which we characterized each of the tiles provided in the first part of this competition using the Reinhard method ^6^, which transforms the RGB color image to the CIELAB colorspace. We then converted the CIELAB projections into the HSV colorspace and then selected 5 representative tiles from the spectrums observed at the 10th, 25th, 50th, 75th, and 90th percentiles for both hue (H) and value (V). We then made 10 color transformation for each input image to the 10 representative tiles identified above using the Reinhard color transfer method, thus constituting our image color augmentation approach.

### Neural Network Architecture

In initial tests using ResNet50 as a pre-trained feature extractor for the pathology image tiles, we noticed a lack of resolution and detail as many of these pre-trained CNN’s take fairly small images (244 ×; 244 pixels). In order to maintain resolution of important features of the pathology while also approaching the problem with the inspiration of how a pathologist examines slides, we decided to convolve ResNet50 at two magnifications (100×100 μm, 400×400 μm) with a stride length that was half of the kernel length (**Figure 1**). The output of each convolution was a [n-windows, 2048] feature map on which we took the maximum value for each ResNet50 feature. This feature extraction was done with the Keras implementation of ResNet50 where the top layer was not included and an average pooling was done to obtain the 2048 features for each window. At the end of this step, each magnification has a 2048-element vector that reflects the presence of a given ResNet50 feature at a given magnification of the pathology image tile. At this point, we concatenate the 2048-element vector from both magnifications used to create a [2,2048] tensor that is then flattened and used as an input to a trainable fully-connected layer of 512 neurons, trained with a 20% dropout rate, followed by the final multi-class classification layer for the 4 output classes (normal, benign, in-situ, and invasive) with a softmax activation layer. Creation/training of this part of the neural network was implemented in tensorflow.

**Figure 1.**
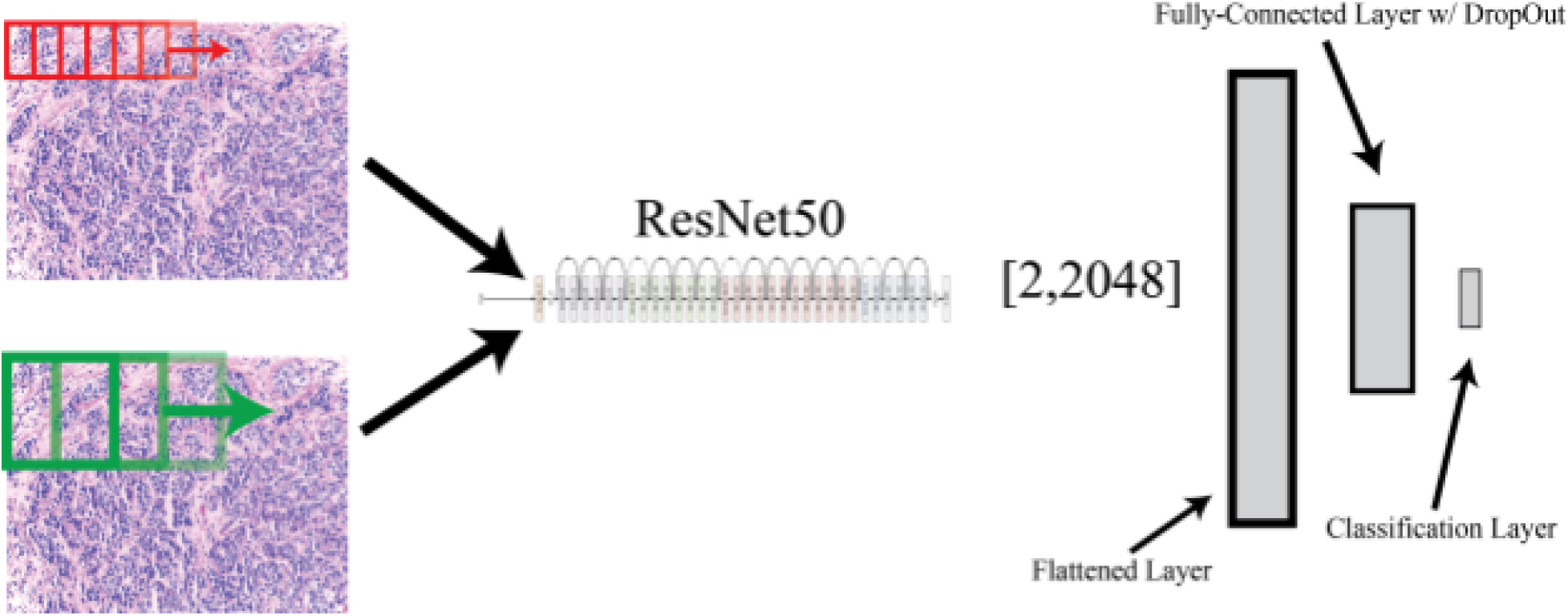
Feature Extraction & Neural Net Architecture. ResNet50 was convolved across a given image tile at two magnifications to generate a ResNet50 feature map for each magnification for each image tile. These feature maps were then flattened into a fully connected layer, on which a secondary fully-connected layer (512 neurons) followed by a final output classification layer wìth a softmax activation.

### Training

In order to train the fully-connected layers of the CNN, we split up our data-set into training, validation, and test sets. We used an 90/5/ split across these respective sets, 5 batches of images per epoch, with 100-fold cross-validation to assess performance via receiver operating characteristic (ROC) curves, confusion matrices to examine individual class accuracy, and measured overall accuracy of the algorithm. We implemented an early stopping approach where after a minimum of 100 epochs, we stopped training when the validation loss did not decrease by 2% in the previous 50 epochs or the validation accuracy did not increase by 5% in the previous 200 epochs. After 100-fold training, we averaged the weights of each training session to arrive at final kernel and bias weights for the fully connected layers of the graph.

## Results

We wanted to assess the ability of neural network to learn the underlying pathologies from two separate sources of pathology images so we conducted a series of experiments where we ran 100x Monte Carlo cross validation with a train/test split of 90/10 percent. We then varied the data available for training, as shown in the figures below while assessing performance on the original Data Set A. We varied the training data based on combination of Data Sets A & B (as described in the methods above) and the use of the image color augmentation approach we implemented, which is described in the methods.

In comparing figure 2 and 3, we observed that the introduction of our image color augmentation approach increased the accuracy in the validation data by a small factor of approximately 2%. In comparing figure 2 and 4, we also noted a small increase of approximately 1 % in performance when including the additional images (Data Set B) derived from the WSI data also provided in the ICIAR 2018 BACH challenge. Somewhat surprisingly, when comparing figure 2 to figure 5, we did not observe an increase in performance with image color augmentation applied to the training set that include data sets A & B.

**Figure 2.**
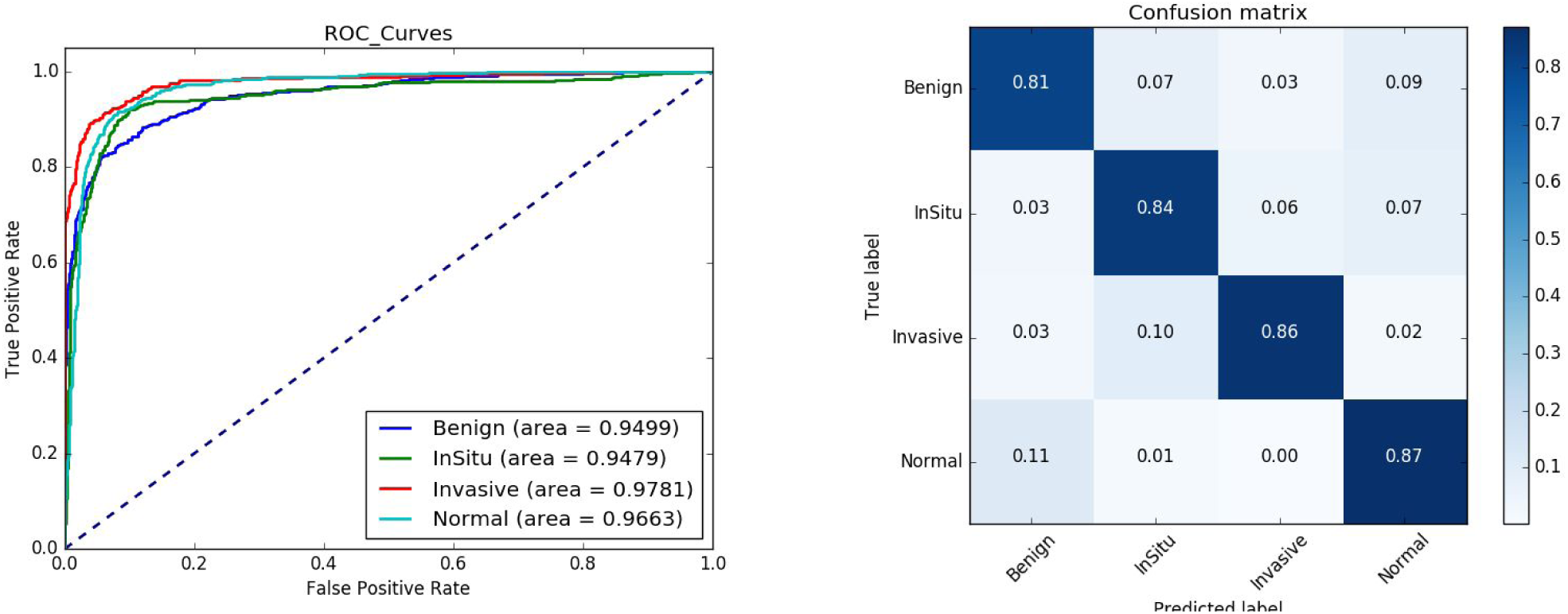
Train on Data Set A, Test on Data Set A - No Image Color Augmentation. With these 400 images, we obtained an overall accuracy of 84.3% across all four classes.

**Figure 3.**
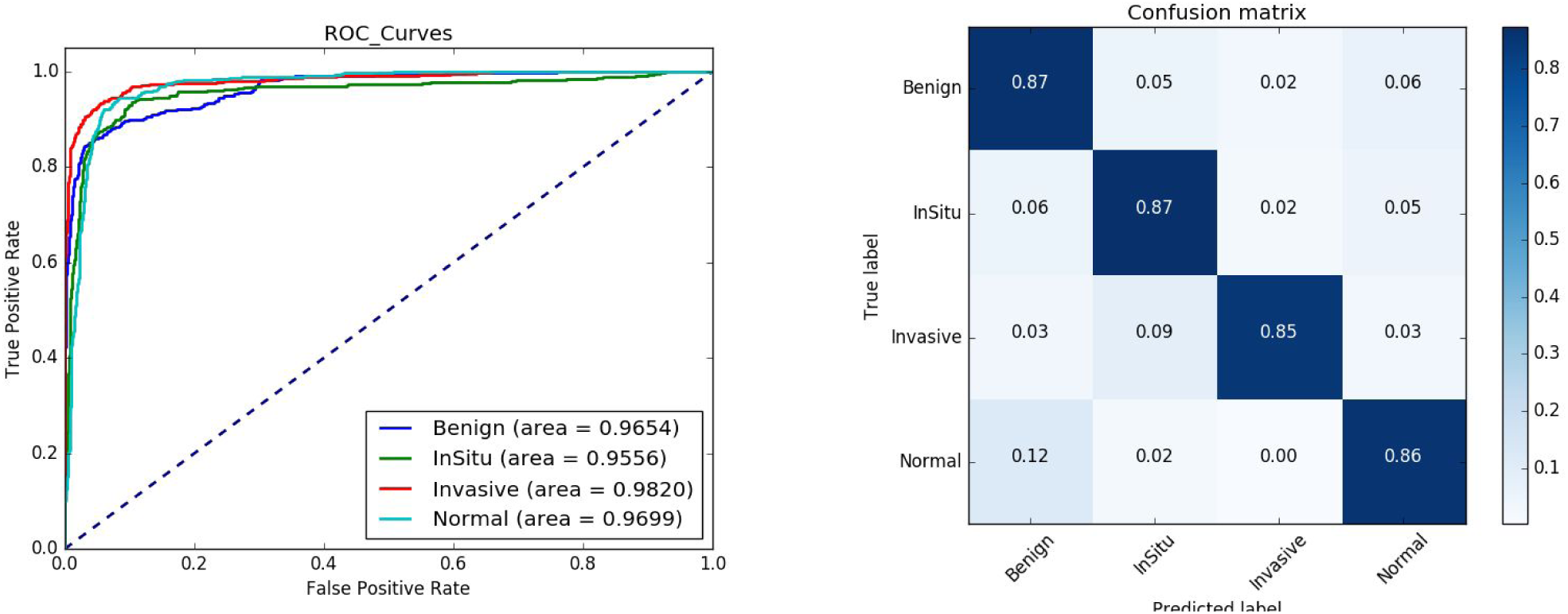
Train on Data Set A, Test on Data Set A - Image Color Augmentation. When introducing augmented images into the training set, we saw an increase in overall accuracy up to 86.2%.

**Figure 4.**
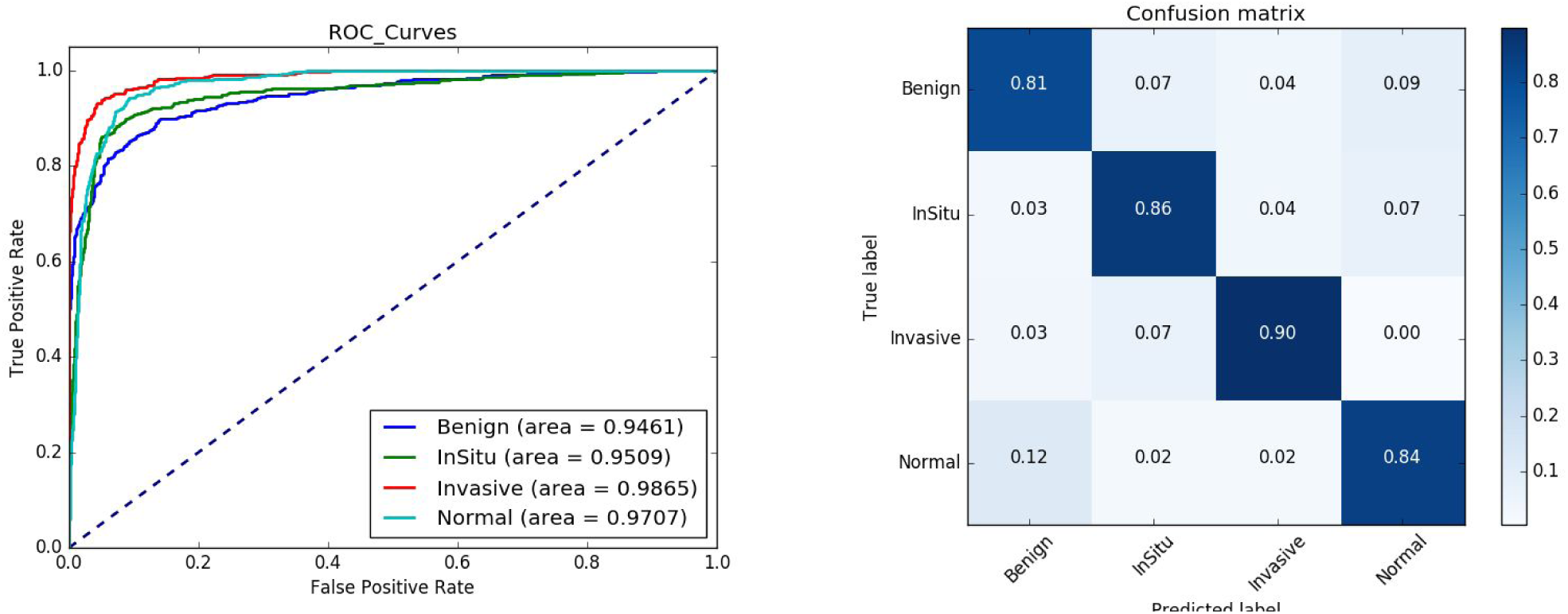
Train on Data Set A & Data Set B, Test on Data Set A - No Image Color Augmentation. When introducing a second data set into our training set, we saw an increase in overall accuracy up to 85.3%.

**Figure 5.**
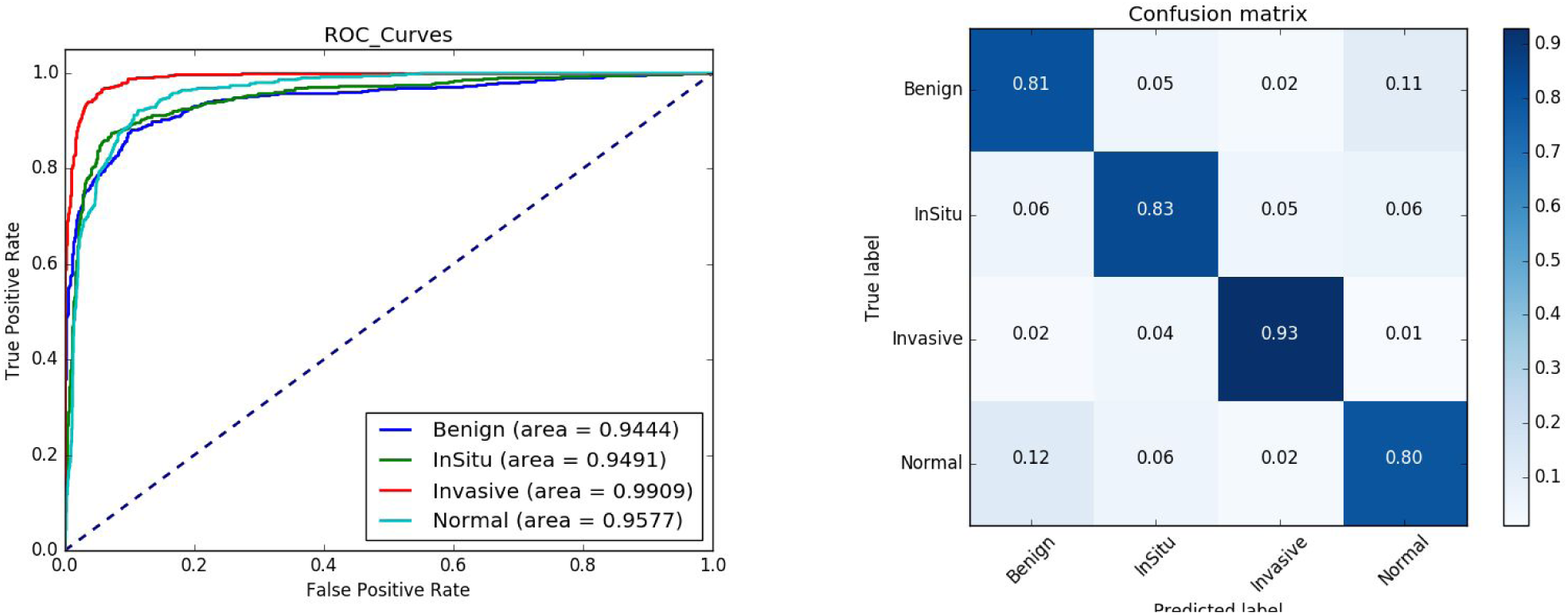
Train on Data Set A & Data Set B, Test on Data Set A - With Image Color Augmentation. When introducing color augmentation into the training of both data sets A and B, we observed an overall accuracy of 84.4%.

## Conclusion

Here we present a method of convoling ResNet50, a pre-trained convolutional neural network, across pathology image tiles at various magnifications to identify low and high-level features that can be later fed into trainable fully-connected layers for the purpose of accurately classifying various types of breast lesions. While we were able to get a respectable accuracy through this method, we believe we were still making mistakes in classification that would be considered obvious to a pathologist. While we never explored the idea of re-training or fine-tuning the weights *within* ResNet50, we hypothesize this approach may be able to push the performance further beyond what we have been able to show here.

Furthermore, when examining the clinical utility of such an algorithm, we felt future directions should focus differentiating cancer from non-cancer primarily as algorithms such as this one will initially have value in the capacity of being screening tests. We believe in order to make this a viable, clinically useful algorithm, future efforts should be placed in ruling out normal images with a high degree of confidence, regardless of the false positive rate for cancer detection by the algorithm.

## References

1. Krizhevsky, A., Sutskever, I. & Hinton, G. E. ImageNet Classification with Deep Convolutional Neural Networks. In Advances in Neural Information Processing Systems 25 (eds. Pereira, F., Burges, C. J. C., Bottou, L. & Weinberger, K. Q.) 1097–1105 (Curran Associates, Inc., 2012).

2. Esteva, A. et al. Dermatologist-level classification of skin cancer with deep neural networks. Nature 542, 115–118 (2017).

3. Djuric, U., Zadeh, G., Aldape, K. & Diamandis, P. Precision histology: how deep learning is poised to revitalize histomorphology for personalized cancer care. npj Precision Oncology 1, 22 (2017).

4. Lee, J.-G. et al. Deep Learning in Medical Imaging: General Overview. Korean J. Radiol. 18, 570–584 (2017).

5. Kao, C.-Y. & McMillan, L. A Novel Deep Learning Architecture for Testis Histology Image Classification. arXiv [cs.CV] (2017).

6. Reinhard, E., Adhikhmin, M., Gooch, B. & Shirley, P. Color transfer between images. IEEE Comput. Graph. Appl. 21, 34–41 (2001).

